# Estrogen Receptor 1 Signaling in Hepatic Stellate Cells Designates Resistance to Liver Fibrosis

**DOI:** 10.1101/2024.10.10.617541

**Authors:** Tianhao Li, Gang Wang, Han Zhao, Fuhai Liu, Dingbao Chen, Xin Zhou, Zhangyuzi Deng, Ying Cao, Wei Fu, Haoyue Zhang, Jing Yang

**Affiliations:** State Key Laboratory of Membrane Biology, School of Life Sciences, Peking University, Beijing, China, 100871; Center for Life Sciences, Academy for Advanced Interdisciplinary Studies, Peking University, Beijing, China, 100871; IDG/McGovern Institute for Brain Research, Peking University, Beijing, China, 100871; Department of Hepatobiliary, Peking University People’s Hospital, Beijing, China, 100044; Department of General Surgery, Peking University Third Hospital, Beijing, China, 100191; Cancer Center, Peking University Third Hospital, Beijing, China, 100191; Institute of Molecular Physiology, Shenzhen Bay Laboratory, Shenzhen, Guangdong, China, 518055

**Keywords:** Liver fibrosis, hepatic stellate cells (HSCs), sex hormone, estrogen receptor 1 (ESR1), sexual dimorphism

## Abstract

The prevalence and severity of liver fibrosis appear higher in men than in premenopausal women, while postmenopausal women exhibit the worsened disease. However, the pathophysiological mechanism underlying such clinical observations remains incompletely understood. Here, we show that sex hormone depletion in adult female mice exaggerates the model of liver fibrosis, while estradiol replacement in castrated male mice is sufficient to mitigate the disease severity. Transcriptomic analyses and immunohistochemistry then demonstrate that both human and mouse hepatic stellate cells (HSCs), the primary cell type responsible for extracellular fibrous depositions, predominantly express the estrogen receptor 1 (ESR1). Of importance, genetic deletion of ESR1 in mouse HSCs markedly promotes liver fibrosis. Moreover, chromatin immunoprecipitation followed by sequencing (ChIP-seq) and in vitro manipulations reveal that ESR1 can directly target the expression of fibrosis-related genes in HSCs. Together, this study has elucidated a critical aspect of ESR1 signaling in the sexual dimorphism of liver fibrosis.

## Introduction

Liver fibrosis is a pathological event defined by the excessive deposition of extracellular matrix proteins, particularly collagens, in this vital organ. This disease condition often accompanies various types of chronic damage to hepatocytes, e.g., viral hepatitis, biliary obstruction, alcoholic liver disease, or nonalcoholic fatty liver disease (NAFLD, or metabolic dysfunction-associated steatohepatitis as recently defined). While the formation of fibrotic scars represents an integrative step of tissue repair, this process can result in the impairment of normal liver functions and eventually progress to more severe complications, i.e., cirrhosis, liver failure, and hepatocellular carcinoma, that significantly impact the morbidity and mortality of patients (Bataller and Brenner, 2005; Hernandez-Gea and Friedman, 2011; Kisseleva and Brenner, 2021; Taylor et al., 2020).

Extensive studies have documented that hepatic stellate cells (HSCs) are the primary cell type involved in the production of extracellular matrix proteins during liver fibrosis (Friedman, 2008; Kamm and McCommis, 2022; Tsuchida and Friedman, 2017; Yin et al., 2013). Under the healthy condition, HSCs are in a quiescent state characterized by the stellate morphology and the storage of vitamin A (i.e., retinol) in their lipid droplets. Upon inflammatory or oxidative stimuli during liver injuries, HSCs are triggered into a proliferative, fibrogenic state with the loss of stellate morphology, decrease of vitamin A storage, and enhanced release of extracellular matrix proteins. It has been broadly recognized that activated HSCs, through their complex crosstalk with Kupffer cells, T lymphocytes, sinusoidal endothelial cells, and other cell types, designate the outcome of liver fibrosis. Therefore, a comprehensive understanding of mechanisms regulating the fibrogenic response of HSCs is essential for effective therapeutic strategies against this debilitating disease (Ezhilarasan et al., 2018; Higashi et al., 2017).

Of importance, clinical evidence has indicated that the prevalence and severity of liver fibrosis exhibit significant gender differences (Ballestri et al., 2017; Buzzetti et al., 2017; Garate-Carrillo et al., 2020; Guy and Peters, 2013; Lefebvre and Staels, 2021; Lonardo et al., 2019; Pan and Fallon, 2014). For instance, in patients with nonalcoholic steatohepatitis (NASH), men were at a higher risk of more severe fibrosis compared to premenopausal women, while postmenopausal women had a similar disease severity compared to men (Ciardullo et al., 2022; Yang et al., 2014). In addition, in patients with chronic Hepatitis B or Hepatitis C infection, females had a delayed progression of liver fibrosis than males, but this protective effect was lost in postmenopausal women (Saif-Al-Islam et al., 2020; Xiong et al., 2019). Those observations have suggested that estrogen may enable the resistance to liver fibrosis. In support of this notion, the incidence of NAFLD was lower in the women taking hormonal replacement therapy compared to postmenopausal women (Hamaguchi et al., 2012). Also, estrogen treatment could mitigate the severity of liver fibrosis and HSC activation in rodent models (Mondal et al., 2023; Shimizu et al., 1999; Yasuda et al., 1999; Zhang et al., 2018). However, the pathophysiological mechanism underlying this estrogen modulation of HSCs remains incompletely charted.

## Results & Discussion

As the entry point of this study, we exploited the procedure of bilateral ovariectomy in adult C57BL/6 wild-type female mice to mimic menopausal hormone depletion (Medina-Contreras et al., 2020) (Fig. 1A). Compared to the mice that underwent sham surgery, the ovariectomized mice fed with the normal chow diet (NCD) did not exhibit liver steatosis or fibrosis in the timeframe of our experimental setup (Fig. 1B and 1C). We then subjected the mice to the methionine-choline deficient (MCD) diet, a standard metabolic model to induce liver fibrosis (Constandinou et al., 2005; Yanguas et al., 2016). Of importance, Sirius Red staining, a classic histochemical method to visualize collagen fibers in tissues, revealed a worsened level of liver fibrosis in the ovariectomized mice challenged by the MCD diet (Fig. 1C and 1D). Also, the immunohistochemical assessment of alpha-smooth muscle actin (α-SMA), a cellular marker for activated HSCs (Carpino et al., 2005), showed its enhanced expression in the livers of MCD-fed mice receiving ovariectomy compared to sham surgery (Fig. 1E and 1F). In addition, it has been documented that activated HSCs express the tissue inhibitor of metalloproteinase 1 (TIMP1) to facilitate the deposition of extracellular matrix by blocking the activity of matrix metalloproteinases (MMPs) (Benyon and Arthur, 2001; Kisseleva and Brenner, 2006). Accordingly, TIMP1 protein levels were markedly increased in the liver tissues of ovariectomized mice fed with the MCD diet as examined by immunofluorescence staining (Fig. 1G and 1H). Moreover, previous studies with single-cell RNA sequencing (scRNA-seq) analyses have identified several collagen genes that are highly expressed by mouse HSCs during liver fibrosis, e.g., *Col1a1*, *Col1a2*, and *Col3a1* (Dobie et al., 2019). Indeed, mRNA levels of those fibrosis-related genes were significantly elevated in the livers of the MCD-fed mice receiving ovariectomy, as determined by quantitative PCR (qPCR) analyses (Fig. 1I). These results confirmed that estrogen deletion in adult female mice could exaggerate liver fibrosis.

**Figure 1.**
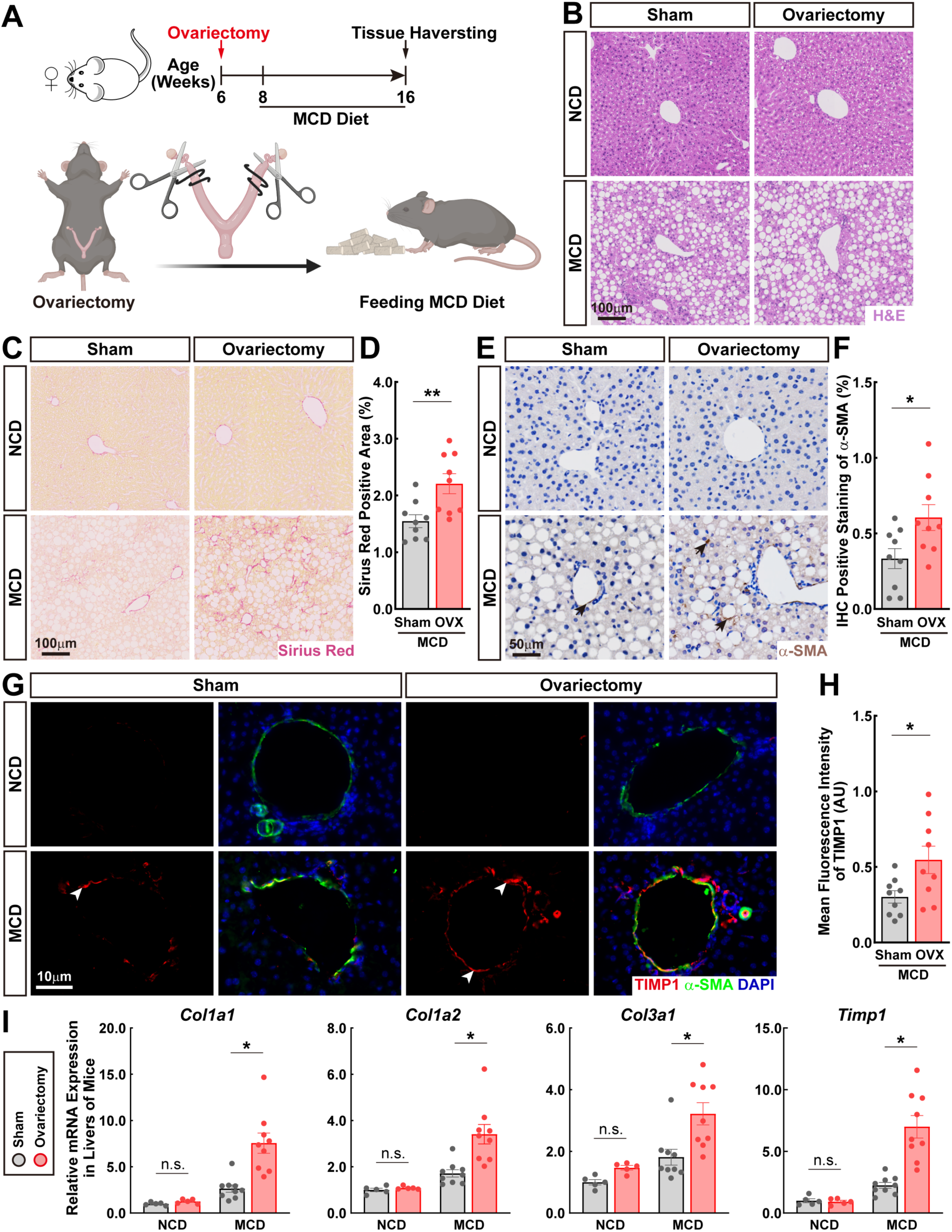
Sex hormone depletion in adult female mice exaggerates the MCD diet-induced liver fibrosis. C57BL/6 wild-type female mice of 6 weeks old underwent ovariectomy or sham surgery. The mice were then fed with the normal chow diet (NCD) or methionine-choline deficient (MCD) diet to induce liver fibrosis. **(A)** Diagram of the experimental procedure. **(B to F)** Paraffin sections of the liver tissues of indicated conditions were assessed by histochemistry or immunohistochemistry. **(B)** Representative images of H&E staining. **(C and D)** Representative images of Sirius Red staining **(C)** and the quantification of the percentage (%) of Sirius Red-positive area **(D)**. **(E and F)** Representative images of anti-α-SMA immunohistochemistry **(E**, black arrows exemplify anti-α-SMA signals) and the quantification of the percentage (%) of α-SMA-positive area **(F)**. mean ± SEM, * *p* < 0.05, ** *p* < 0.01 (Student’s *t*-test). **(G and H)** Cryosections of the liver tissues were examined by the immunofluorescence co-staining of TIMP1 and α-SMA. **(G)** Representative images were shown. White arrowheads exemplify anti-TIMP1 signals. **(H)** The mean fluorescence intensity of anti-TIMP1 signals was quantified. mean ± SEM, * *p* < 0.05 (Student’s *t*-test). **(I)** mRNA levels of fibrosis-related genes were determined by the qPCR analyses. mean ± SEM, * *p* < 0.05, n.s., not significant (two-way ANOVA test).

We next tested whether estrogen would be sufficient to enact the resistance to liver fibrosis. To this end, we utilized the procedure of bilateral castration and 17β-estradiol hormone replacement in adult C57BL/6 wild-type male mice (Fig. 2A). Importantly, this approach of hormone replacement under the NCD diet condition did not cause liver steatosis or fibrosis in the experimental timeframe (Fig. 2B and 2C). However, the 17β-estradiol treatment strongly suppressed the deposition of collagen fibers in the liver tissues under the MCD diet challenge, as elucidated by Sirius Red staining (Fig. 2C and 2D). Also, expression levels of α-SMA were reduced in the livers of castrated male mice receiving 17β-estradiol replacement (Fig. 2E and 2F). In addition, TIMP1 expression was inhibited in response to the 17β-estradiol treatment of MCD-fed mice (Fig. 2G and 2H). Similarly, mRNA levels of key fibrosis-related genes significantly decreased in the liver tissues of MCD-fed mice receiving 17β-estradiol (Fig. 2I). These results demonstrated that estrogen signaling in castrated adult male mice is sufficient to mitigate liver fibrosis.

**Figure 2.**
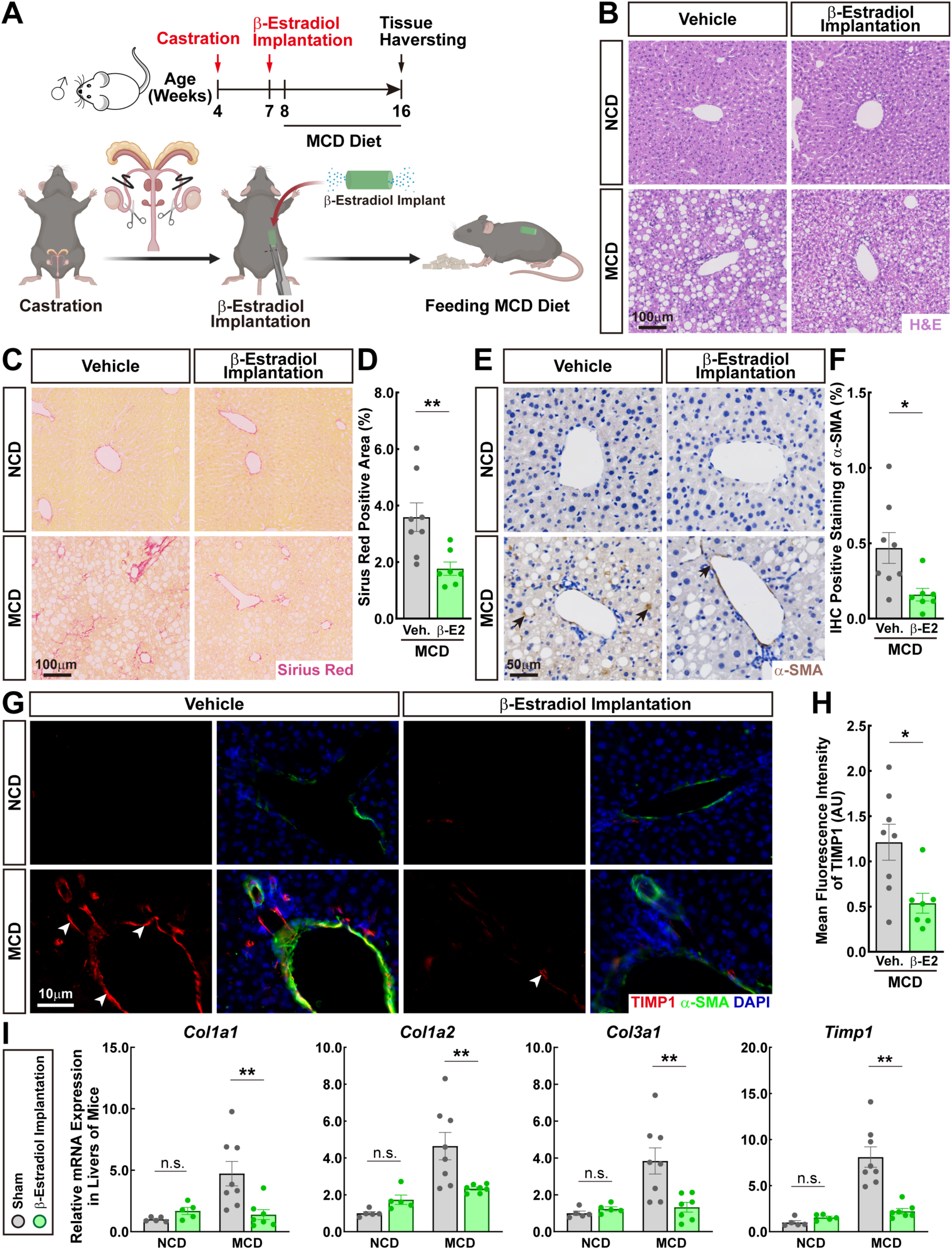
Estrogen replacement in castrated adult male mice is sufficient to mitigate liver fibrosis. C57BL/6 wild-type male mice of 4 weeks old underwent castration and 17β-estradiol (β-E2) hormone replacement. The mice then received the normal chow diet (NCD) or methionine-choline deficient (MCD) diet to induce liver fibrosis. **(A)** Diagram of the experimental procedure. **(B to F)** Paraffin sections of the liver tissues of indicated conditions were assessed by histochemistry or immunohistochemistry. **(B)** Representative images of H&E staining. **(C and D)** Representative images of Sirius Red staining **(C)** and the quantification of the percentage (%) of Sirius Red-positive area **(D)**. **(E and F)** Representative images of anti-α-SMA immunohistochemistry **(E**, black arrows exemplify anti-α-SMA signals) and the quantification of the percentage (%) of α-SMA-positive area **(F)**. mean ± SEM, * *p* < 0.05, ** *p* < 0.01 (Student’s *t*-test). **(G and H)** Cryosections of the liver tissues were examined by the immunofluorescence co-staining of TIMP1 and α-SMA. **(G)** Representative images were shown. White arrowheads exemplify anti-TIMP1 signals. **(H)** The mean fluorescence intensity of anti-TIMP1 signals was quantified. mean ± SEM, * *p* < 0.05 (Student’s *t*-test). **(I)** mRNA levels of fibrosis-related genes were determined by the qPCR analyses. mean ± SEM, ** *p* < 0.01, n.s., not significant (two-way ANOVA test).

We sought to determine the molecular mechanism of estrogen signaling in HSCs. HSCs were isolated from the non-parenchymal cells of human liver tissues by fluorescence-activated cell sorting (FACS), taking advantage of the autofluorescence property of retinol at 405nm (Mederacke et al., 2015). Human HSCs, which were defined as retinol^+^ CD45^-^ (Fig. 3A), were then profiled by RNA sequencing (RNA-seq). Notably, based on conventional methods such as reverse transcriptase PCR, it was repeatedly reported that rodent HSCs only express the estrogen receptor 2 (ESR2, also known as ERβ) for estrogen signaling (McCarty et al., 2009; Mondal et al., 2023; Que et al., 2018; Zhang et al., 2018; Zhou et al., 2001). However, the validity of this canonical view in the field is untested with more advanced transcriptomic technologies. We observed that HSCs from both male and female human livers predominantly express *ESR1* (also known as ERα) but not *ESR2* (Fig. 3C). Also, recent studies suggested that human HSCs might express the G protein-coupled estrogen receptor (GPER) (Chen et al., 2021; Cortes et al., 2019). However, the RNA-seq data exhibited a relatively minor level of *GPER* in male or female human HSCs (Fig. 3C). On the other hand, mRNA levels of androgen receptor (AR) appeared undetectable in human HSCs (Fig. 3C). We confirmed this specific expression of ESR1 in human HSCs by immunofluorescence co-staining of ESR1 and α-SMA, which unequivocally identified the ESR1-positive nuclei of α-SMA-positive HSCs in male or female human livers (Fig. 3D). In contrast, while anti-ESR2 immunofluorescence staining revealed the ESR2-positive cells in human liver tissues, they did not overlap with α-SMA-positive HSCs (Fig. 3E).

In parallel, we FACS-sorted retinol^+^ CD45^-^ mouse HSCs from adult C57BL/6 wild-type mice (Fig. 3B). RNA-seq profiling showed that male or female mouse HSCs express *ESR1* but not *ESR2* or *GPER* (Fig. 3C), highly reminiscent of the above observation with human HSCs. Also, we analyzed the published scRNA-seq datasets GSE136103 (Duan et al., 2021) and GSE137720 (Dobie et al., 2019) that contain the non-parenchymal cells of adult C57BL/6 wild-type male mice under the healthy or fibrotic conditions. Within those pooled datasets, mouse HSCs could be clearly distinguished from other cell types, e.g., vascular smooth muscle cells (VSMC) and fibroblasts (FB) (Fig. 3F). Of importance, at this single-cell resolution, mouse HSCs primarily expressed *ESR1* in both healthy and fibrotic conditions (Fig. 3G), in accordance with the conclusion obtained from our RNA-seq data.

**Figure 3.**
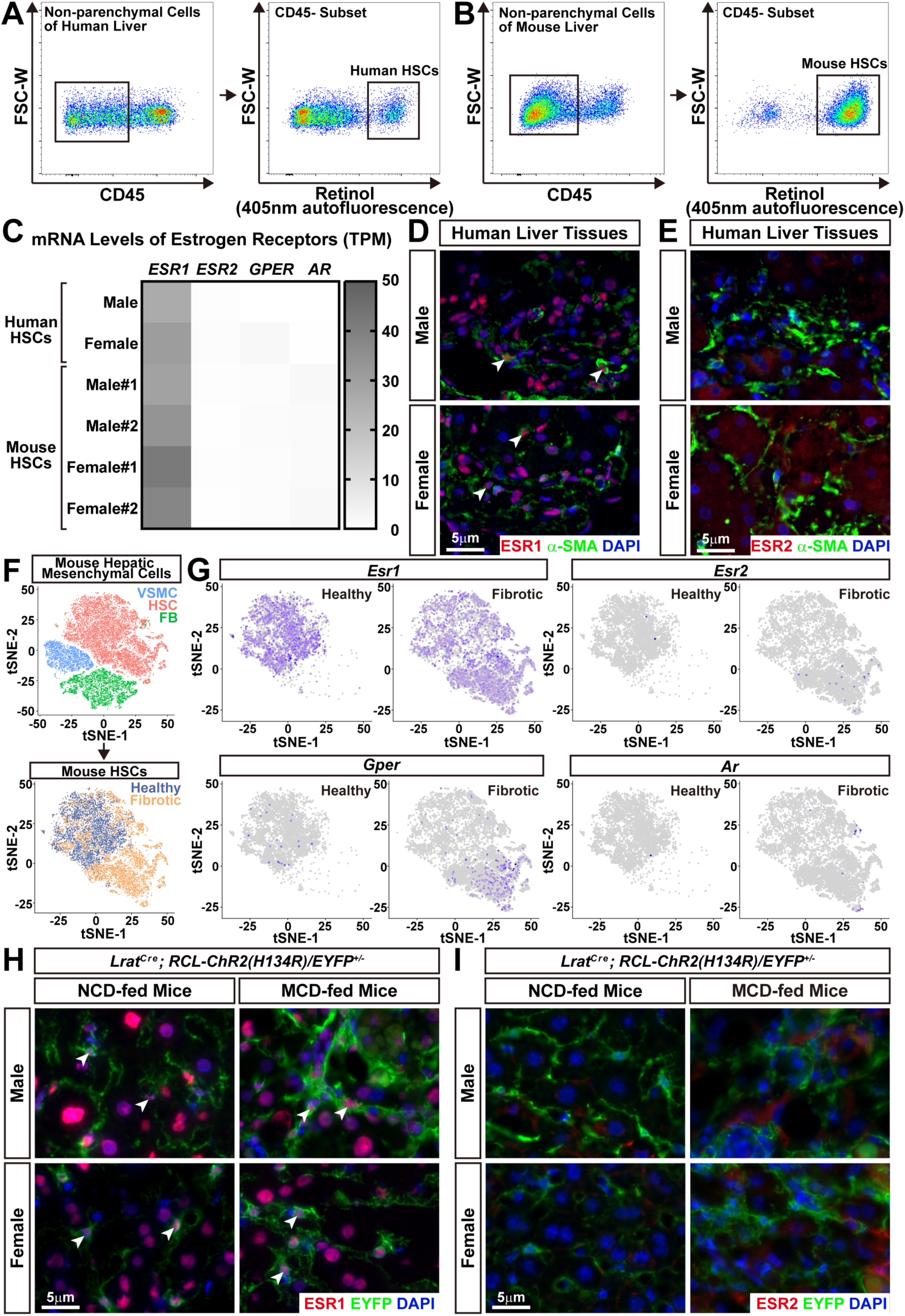
Human and mouse HSCs predominantly express ESR1. **(A to C)** Human or mouse HSCs were FACS-sorted and subjected to RNA-seq. **(A and B)** Representative FACS plots of human HSCs **(A)** or C57BL/6 wild-type mouse HSCs **(B)** that were identified as retinol (405nm autofluorescence)^+^ CD45^-^. **(C)** mRNA levels of estrogen receptors *ESR1* and *ESR2* in human HSCs (from male or female patients) and mouse HSCs (from male or female mice; two replicates for each gender) were determined by RNA-seq. **(D and E)** Paraffin sections of the human liver tissues of male or female patients were examined by the immunofluorescence co-staining of α-SMA with ESR1 **(D)** or ESR2 **(E)**. White arrowheads exemplify the ESR1-positive nuclei of α-SMA-positive HSCs. **(F and G)** The published scRNA-seq datasets GSE136103 and GSE137720 of the non-parenchymal cells of adult C57BL/6 wild-type male mice were analyzed. **(F)** t-Distributed stochastic neighbor embedding (t-SNE) plots of the pool datasets (upper panel) and mouse HSCs identified in the control healthy condition or the model of liver fibrosis (lower panel). VSMC, vascular smooth muscle cells; FB, fibroblasts. **(G)** Feature plots of the indicated genes in mouse HSCs of the healthy and fibrotic conditions. **(H and I)** *Lrat^Cre^; RCL-ChR2(H134R)/EYFP^+/-^* male or female mice of 8 weeks old were fed with the normal chow diet (NCD) or methionine-choline deficient (MCD) diet to induce liver fibrosis. Cryosections of the liver tissues were examined by the immunofluorescence co-staining of EYFP with ESR1 **(F)** or ESR2 **(G)**. White arrowheads exemplify the ESR1-positive nuclei of EYFP-positive HSCs.

We noted that HSCs decrease their retinol storage upon activation (Blaner et al., 2009; Kamm and McCommis, 2022; Tsuchida and Friedman, 2017), which may pose an intrinsic limitation of the FACS-based approach for HSC isolation. We thus circumvented this issue by exploiting the *Lrat^Cre^* mouse line that specifically drives the Cre recombinase in HSCs. Consistent with the previous documentation (Mederacke et al., 2013), >99% of HSCs were labeled as EYFP-positive in the livers of *Lrat^Cre^; RCL-ChR2(H134R)/EYFP^+/-^* reporter mice (Fig. 4A). *Lrat^Cre^; RCL-ChR2(H134R)/EYFP^+/-^* male or female mice were then fed with the NCD or the MCD diet to induce liver fibrosis. Immunofluorescence co-staining of ESR1 and EYFP evidently identified the ESR1-positive nuclei of EYFP-positive HSCs in the male or female liver tissues of NCD-fed or MCD-fed conditions (Fig. 3H). On the contrary, there were no detectable ESR2-positive HSCs in all the examined conditions (Fig. 3I). Together, these results established that for both genders, human and mouse HSCs predominantly express ESR1.

**Figure 4.**
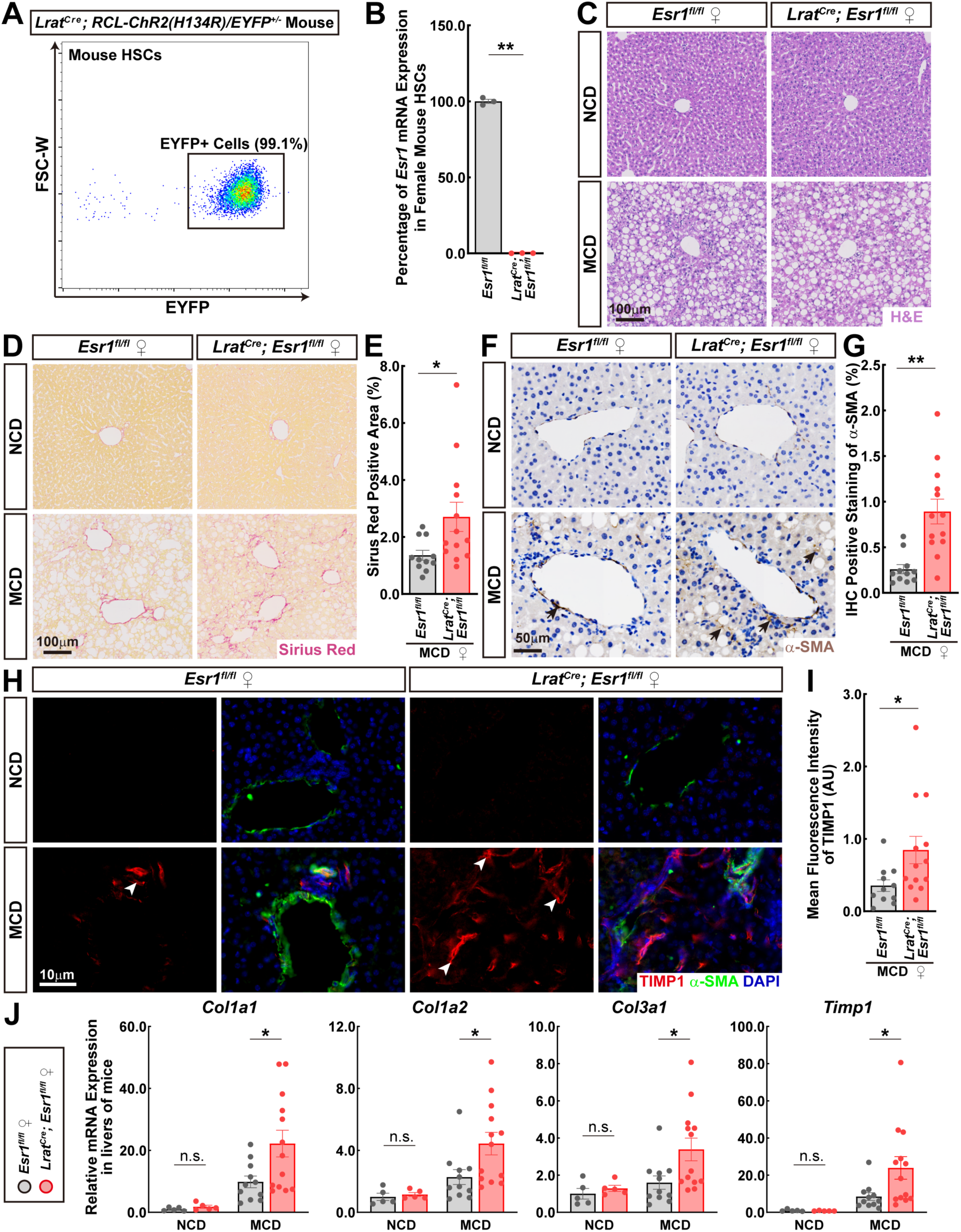
Genetic blockage of ESR1 signaling in HSCs aggregates the model of liver fibrosis. **(A)** HSCs in the liver tissues of *Lrat^Cre^; RCL-ChR2(H134R)/EYFP^+/-^* mice fed with the normal chow diet were examined by FACS. A vast majority (>99%) of retinol^+^ CD45^-^ HSCs were EYFP-positive. **(B to J)** *Lrat^Cre^;Esr1^fl/fl^* or control *Esr1^fl/fl^* female littermates of 8 weeks old received the normal chow diet (NCD) or methionine-choline deficient (MCD) diet to induce liver fibrosis. **(B)** HSCs were FACS-sorted from the liver tissues of *Lrat^Cre^;Esr1^fl/fl^* or control *Esr1^fl/fl^* mice. Genetic deletion of *Esr1* in *Lrat^Cre^;Esr1^fl/fl^* HSCs was validated by the qPCR analysis. mean ± SEM, ** *p* < 0.01 (Student’s *t*-test). **(C to G)** Paraffin sections of the liver tissues of indicated conditions were assessed by histochemistry or immunohistochemistry. **(C)** Representative images of H&E staining. **(D and E)** Representative images of Sirius Red staining **(D)** and the quantification of the percentage (%) of Sirius Red-positive area **(E)**. **(F and G)** Representative images of anti-α-SMA immunohistochemistry **(F**, black arrows exemplify anti-α-SMA signals) and the quantification of the percentage (%) of α-SMA-positive area **(G)**. mean ± SEM, * *p* < 0.05, ** *p* < 0.01 (Student’s *t*-test). **(H and I)** Cryosections of the liver tissues were examined by the immunofluorescence co-staining of TIMP1 and α-SMA. **(H)** Representative images were shown. White arrowheads exemplify anti-TIMP1 signals. **(HI** The mean fluorescence intensity of anti-TIMP1 signals was quantified. mean ± SEM, * *p* < 0.05 (Student’s *t*-test). **(I)** mRNA levels of fibrosis-related genes were determined by the qPCR analyses. mean ± SEM, * *p* < 0.05, n.s., not significant (two-way ANOVA test).

We went on to examine the functional relevance of ESR1 signaling in HSCs. *Lrat^Cre^;Esr1^fl/fl^* mice were generated, which enabled the specific deletion of ESR1 in HSCs. We validated the efficiency of this genetic approach by the qPCR analysis, showing the complete loss of *Esr1* mRNAs in the HSCs of *Lrat^Cre^;Esr1^fl/fl^*compared to control *Esr1^fl/fl^* female littermates (Fig. 4B). Notably, this genetic blockage of ESR1 signaling would not trigger liver steatosis or fibrosis in the NCD-fed female mice (Fig. 4C and 4D). However, the deposition of collagen fibers in the liver under the MCD diet challenge, as visualized by Sirius Red staining, became worsened in *Lrat^Cre^;Esr1^fl/fl^* female mice (Fig. 4D and 4E). Meanwhile, protein levels of α-SMA (Fig. 4F and 4G) or TIMP1 (Fig. 4H and 4I) were significantly elevated in the liver tissues of MCD-fed *Lrat^Cre^;Esr1^fl/fl^* female mice. In addition, mRNA levels of fibrosis-related genes markedly increased in the MCD-challenged livers of *Lrat^Cre^;Esr1^fl/fl^*compared to control female littermates (Fig. 4J). These results supported the notion that ESR1 signaling in HSCs exerts a protective role against liver fibrosis.

Finally, we looked into the molecular mechanism of ESR1 signaling in HSCs. It has been well established that upon the estrogen ligand engagement, ESR1 can regulate the expression of target genes whose promoters or enhancer regions contain the specific sequence of estrogen-responsive element (ERE). Therefore, we systemically profiled the ESR1-binding sites in mouse HSCs by chromatin immunoprecipitation followed by sequencing (ChIP-seq). EYFP-positive HSCs were FACS-sorted from the *Lrat^Cre^; RCL-ChR2(H134R)/EYFP^+/-^* female mice subjected to liver fibrosis and then processed for anti-ESR1 ChIP-seq analyses (Fig. 5A). Approximately a half of identified ESR1-binding sites in the genome of mouse HSCs were located in gene regions including the promoter, untranslated regions (UTRs), exons, and introns, while the other half of ESR1-binding sites resided in distal intergenic regions (Fig. 5B). As expected, *de novo* motif enrichment analysis showed that ERE was the most significantly enriched motif within those ESR1-binding sites, validating the success of anti-ESR1 ChIP-seq procedure (Fig. 5C). Notably, the binding motifs of other nuclear receptors were also identified (Fig. 5C), e.g., nuclear receptor subfamily 2 group F member 6 (NR2F6, also known as EAR2) and nuclear receptor subfamily 2 group F member 2 (NR2F2, also known as COUP-TFII). This observation may implicate the potential genome-wide cooperative action of those nuclear receptors with ESR1. Of importance, we identified the ESR1-binding sites at the loci of key fibrosis-related genes, e.g., *Col1a1*, *Col1a2*, *Col3a1*, and *Timp1* (Fig. 5D), suggesting a direct transcriptional control by ESR1. Indeed, the 17β-estradiol treatment of in vitro cultured mouse HSCs could effectively suppress the mRNA levels of those genes, as assessed by RNA-seq profiling (Fig. 5E). These results demonstrated that ESR1 directly targets the expression of fibrosis-related genes in mouse HSCs.

**Figure 5.**
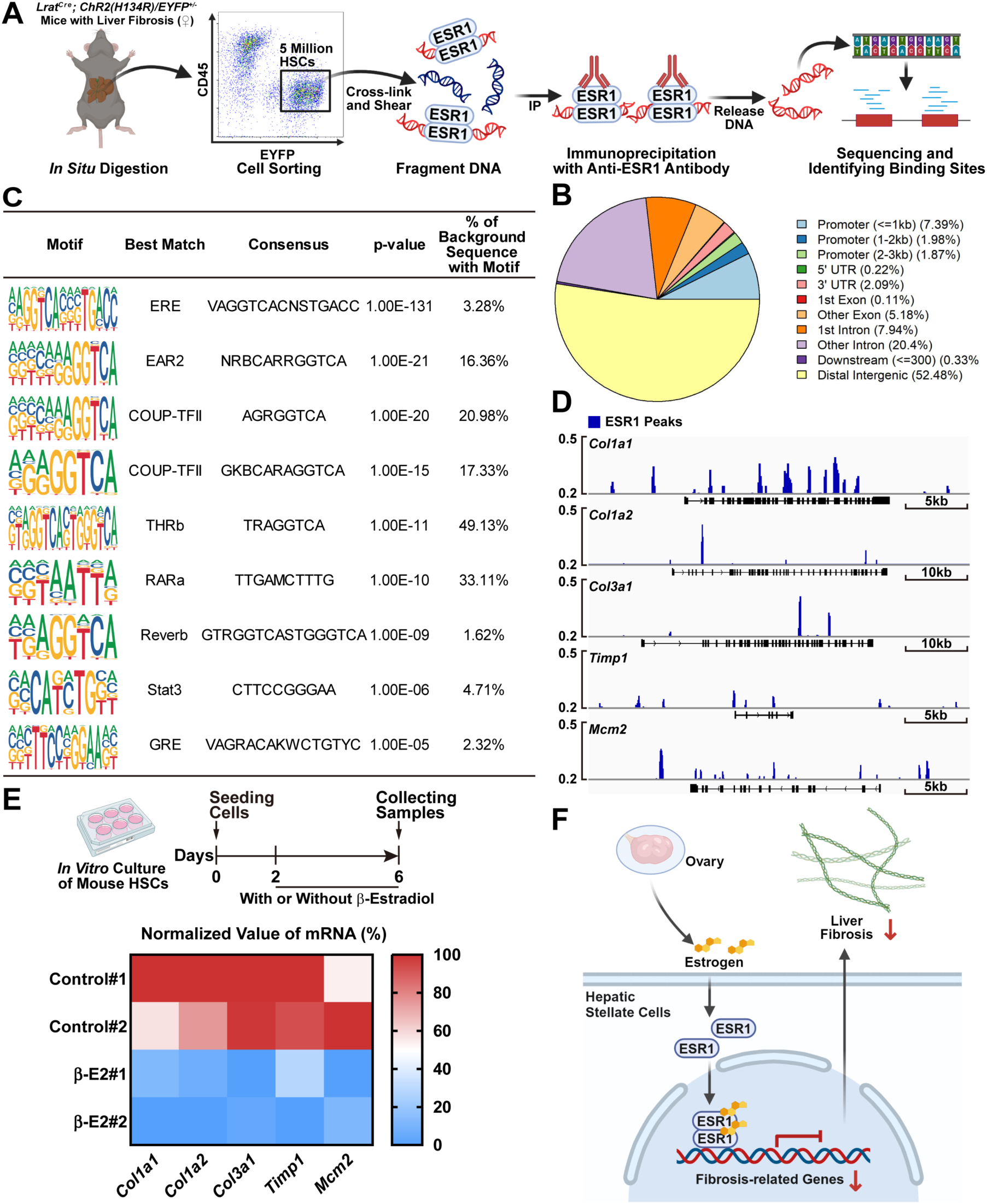
ESR1 directly targets the expression of fibrosis-related genes in mouse HSCs. **(A to D)** *Lrat^Cre^; RCL-ChR2(H134R)/EYFP^+/-^* female mice of 8 weeks old were subjected to the model of liver fibrosis. HSCs were then FACS-sorted from the liver tissues and subjected to anti-ESR1 chromatin immunoprecipitation followed by sequencing (ChIP-seq). **(A)** Diagram of the experimental procedure. **(B)** Summary of the genomic distribution of ESR1-binding sites in mouse HSCs. **(C)** *De novo* motif enrichment analysis showing the motifs identified in ESR1-binding sites in mouse HSCs. **(D)** Genomic browser tracks showing ESR1-binding sites at the loci of fibrosis-related genes in mouse HSCs. **(E)** HSCs were FACS-sorted from the liver tissues of C57BL/6 wild-type female mice for *in vitro* cultures. Mouse HSCs were then treated with 17β-estradiol (β-E2) or vehicle control, and expression levels of fibrosis-related genes were profiled by RNA-seq (two replicates for each condition). **(F)** Diagram of ESR1 signaling in HSCs to limit liver fibrosis.

In sum, we have elucidated a critical aspect of ESR1 signaling in HSCs designating resistance to liver fibrosis (Fig. 5F). A previous study showed that a *cis*-acting regulatory variation in the enhancer region of *ESR1* gene would be associated with the susceptibility of patients to Hepatitis B virus-related liver cirrhosis (Yan et al., 2011), supporting the essential role of ESR1 in the disease context. Also, it was recently reported that ESR1 could directly control the expression of the patatin-like phospholipase domain-containing 3 (PNPLA3) I148M variant in human hepatocytes, thus influencing the women’s susceptibility to NAFLD (Cherubini et al., 2023). In light of our current work, whether this ESR1-PNPLA3 signaling event may also act in HSCs warrants research attention. A comprehensive knowledge of the sexual dimorphism of liver fibrosis and its underlying mechanisms will lead to better stratification of patients for diagnostic or therapeutic strategies.

At the same time, HSCs may be subjected to the impact of other hormones besides estrogen. For instance, genetic or pharmacological blockage of 11β-hydroxysteroid dehydrogenase type 1, the key enzyme that catalyzes the intracellular conversion of cortisone to physiologically active cortisol, ameliorated the mouse models of liver fibrosis, suggesting the critical role of the hormone cortisol in this disease scenario (Lee et al., 2022). Also, deficiency of thyroid hormones is significantly associated with increased risk of liver fibrosis (Piantanida et al., 2020; Ritter et al., 2020; Sinha et al., 2018), and the thyroid hormone receptor agonist has recently become the first FDA-approved treatment of liver fibrosis in patients with NASH (Harrison et al., 2024). Whether those hormones may regulate the fibrogenic state of HSCs calls for in-depth studies. Moreover, recent works by colleagues and us have revealed the dense presence of sympathetic innervations in human and mouse livers (Adori et al., 2021; Liu et al., 2021). Whether HSCs may be influenced by such local neural signals, in addition to those hormones in the blood circulation, appears to be a tempting possibility that awaits future research.

## Materials and Methods

Further information and requests for resources and reagents should be directed to and will be fulfilled by Jing Yang.

### Human liver tissues

Human liver tissues were collected in compliance with the protocols approved by the Institutional Ethics Committee of Peking University People’s Hospital or Peking University Third Hospital, and informed consent was signed by each involved patient. Normal liver tissues were obtained from patients (males and females, 33 ∼ 76 years old, without the detectable infection of Hepatitis B or Hepatitis C virus) during the hepatectomy for benign or malignant tumors.

For the fluorescence-activated cell sorting (FACS) of human hepatic stellate cells (HSCs), the procedure was modified from the published protocol (Mederacke et al., 2015). Fresh human liver tissues were immediately perfused through main blood vessels with 20ml of the Perfusion Solution (99.6mg/l NaH_2_PO_4_·2H_2_O, 242.15mg/l Na_2_HPO_4_·12H_2_O, 350mg/l NaHCO_3_, 2.38g/l HEPES, 8.0g/l NaCl, 400mg/l KCl, 900mg/l glucose, 190mg/l EGTA, pH 7.35∼7.45), 30ml of the Digestion Solution (99.6mg/l NaH_2_PO_4_·2H_2_O, 242.15mg/l Na_2_HPO_4_·12H_2_O, 350mg/l NaHCO_3_, 2.38g/l HEPES, 8.0g/l NaCl, 400mg/l KCl, 560mg/l CaCl_2_·2H_2_O, pH 7.35∼7.45, pre-warmed to 37^°^C) containing 0.5mg/ml pronase (Sigma, #P5147), and 40ml of the Digestion Solution containing 0.625mg/ml collagenase Ⅱ (Biosharp, #BS164). The liver tissues were then minced into small pieces and incubated at 37^°^C for 15 min in the Digestion Solution containing 0.625mg/ml collagenase Ⅱ and 75U/ml DNaseΙ (Sigma, #D5025). The digested samples were mashed through a 70-μm cell strainer and centrifuged at 50*g* for 3min at 4^°^C to remove tissue debris and hepatocytes, and the supernatant containing non-parenchymal cells was collected and further centrifuged at 580*g* for 10min at 4^°^C. The cell pellets were washed in the Gey’s Balanced Salt Solution (GBSS; 121.92mg/l Na_2_HPO_4_·12H_2_O, 30mg/l KH_2_PO_4_, 227mg/l NaHCO_3_, 8.0g/l NaCl, 370mg/l KCl, 210mg/l MgCl_2_·6H_2_O, 70mg/l MgSO_4_·7H_2_O, 225mg/l CaCl_2_·2H_2_O, 991mg/l glucose, pH 7.35∼7.45) and centrifuged again at 580*g* for 10min at 4^°^C. The cell pellets were resuspended in GBSS containing a final concentration of 10% (w/v) Nycodenz (Serumwerk Bernburg AG, #18003). 10ml of cell suspension was loaded into 15ml Falcon tubes and gently overlaid with 1.5 ml GBSS. The cell preparations were centrifuged at 1,380*g* for 17min at 4^°^C. The resulting cell layers between the two fractions were collected and stained with FITC-conjugated anti-human CD45 (BioLegend, #304054) and 7-AAD (Biolegend). The stained non-parenchymal cells of human liver tissues were sorted on BD Aria Fusion, and HSCs were identified as retinol (405nm autofluorescence)^+^ CD45^-^. Total RNAs of FACS-sorted human HSCs were extracted by the RNeasy Mini Kit (Qiagen) and subjected to single-end RNA-seq by the Beijing Genomics Institute. Gene expression levels were normalized as transcripts per million (TPM). The bulk RNA-seq data was deposited to the Sequence Read Archive (https://www.ncbi.nlm.nih.gov/sra) with the accession number GSE275942.

For the immunofluorescence staining, human liver tissues were immediately fixed in phosphate-buffered saline (PBS) containing 3.7% (w/v) paraformaldehyde (PFA) at room temperature for 4h. Liver tissues were then dehydrated in 75% ethanol (diluted in H_2_O) for 2h, 85% ethanol for 2h, 90% ethanol for 2h, 100% ethanol for 45min, a mixture (v:v = 1:1) of ethanol and dimethylbenzene (Sinopharm, #10023418) for 8min, dimethylbenzene for 8min twice, and paraffin for 1h three times. The paraffin-embedded tissues were sectioned at 4-μm thickness. The sections were rehydrated in dimethylbenzene for 15min twice, 100% ethanol for 5min twice, 75% ethanol for 5min, and sterile water for 5min twice. The sections were then incubated in citrate antigen retrieval solution (Solarbio, #C1032) at 95^°^C for 8min, followed by 3% (w/v) hydrogen peroxide solution for 20min. The sections were blocked with PBS containing 3% bovine serum albumin (BSA; Solarbio, #A8010) and immunostained with the intended primary antibodies, including rabbit anti-ESR1 antibody (Millipore, #06-935), rabbit anti-ESR2 antibody (Invitrogen, #PA1311), and goat anti-α-SMA antibody (Novus, #NB300-978). The sections were finally stained with the corresponding Alexa Fluor-conjugated secondary antibodies, and fluorescence images were scanned by Olympus SLIDEVIEW VS200.

### Mouse information and procedures

All the experimental procedures in mice were performed in compliance with the protocol approved by the Institutional Animal Care and Use Committee (IACUC) of Peking University. Mice were maintained on the 12-hr/12-hr light/dark cycle (light period 7:00 am ∼ 7:00 pm), with the chow diet and water available *ad libitum* unless otherwise specified. The mice used in the experiments were 8 ∼ 10 weeks old unless otherwise specified, and the gender of mice for each experiment was specified in figure legends. C57BL/6 wild-type mice were purchased from Charles River International. *Lrat^Cre^* (Cyagen, #C001205), *Esr1^fl/fl^* (Cyagen, #S-CKO-02261), and *Ai32*/*RCL-ChR2(H134R)/EYFP^+/+^* (Jackson Laboratory, #012569) were purchased and in-house bred to generate *Lrat^Cre^;Esr1^fl/fl^* or *Lrat^Cre^; RCL-ChR2(H134R)/EYFP^+/-^* mouse lines.

For the model of liver fibrosis, the mice of indicated conditions were fed with methionine-choline deficient (MCD) diet (Medicience, #MD12052) for 8 weeks.

For the bilateral ovariectomy, C57BL/6 wild-type female mice of 6 weeks old were anesthetized with 3% isoflurane, and the back skin was shaved and prepared with iodine and alcohol. The skin and peritoneum incisions were made below the 13^th^-rib level on both sides to expose ovaries in the peritoneal cavity. The oviduct and blood vessels supplying each ovary were ligated with a 5/0 suture thread before the ovaries on both sides were resected. Finally, the incisions on the peritoneum and the skin were sutured. Sham surgery included all the steps except the ligation and removal of ovaries. The mice were then subjected to the MCD diet treatment 2 weeks after ovariectomy.

For the castration and 17β-estradiol hormone replacement, C57BL/6 wild-type male mice of 4 weeks old were anesthetized with 3% isoflurane. The skin region of the scrotum was shaved and prepared with iodine and alcohol, and a midline incision was made. The testes, vas deferens, and the attached connective tissues on both sides were pulled out. After ligating the vas deferens and blood vessels supplying each testis with a 5/0 suture thread, the testes on both sides were removed. The skin incision was then closed by wound clips (Fine Science Tools). At 3 weeks post-castration, the mice were anesthetized again with 3% isoflurane, and the back skin was shaved and prepared with iodine and alcohol. An incision was made along the skin region of the lower left back, and a 17β-estradiol extended-release pellet (Shinnobio, #XLME-90-B; hormone delivery of 3μg per day) was subcutaneously implanted for each mouse. Finally, the skin incision was closed by a suture. Sham surgery included all the steps except the implantation of 17β-estradiol pellets. The mice were then subjected to the MCD diet treatment 1 week after receiving the hormone replacement.

### Histology of mouse liver tissues

The mice of indicated conditions were perfused with 20ml PBS, followed by 20 ml PBS containing 3.7% (w/v) PFA. Liver tissues were dissected and post-fixed in PBS / 3.7% PFA at room temperature for 4h. For the cryosectioning, mouse liver tissues were preserved in PBS containing 30% (w/v) sucrose at 4^°^C for 24h and then embedded in the optimal cutting temperature compound (OCT; Tissue-Tek, #4583). For the immunofluorescence staining, 10-μm cryosections were blocked with PBS / 3% BSA and immunostained with the intended primary antibodies, including rabbit anti-ESR1 antibody (Millipore, #06-935), rabbit anti-ESR2 antibody (Invitrogen, #PA1311), goat anti-α-SMA antibody (Novus, #NB300-978), rabbit anti-TIMP1 antibody (Bioss, #bs-0415R), and chicken anti-GFP antibody (Aves Labs, #GFP-1010). The sections were then stained with the corresponding Alexa Fluor-conjugated secondary antibodies, and fluorescence images were scanned by Olympus SLIDEVIEW VS200. The mean fluorescence intensity of anti-TIMP1 immunofluorescence signals was quantified in ImageJ (https://imagej.net/ij).

For the paraffin sectioning, mouse liver tissues were dehydrated in 75% ethanol for 2h, 85% ethanol for 2h, 90% ethanol for 2h, 100% ethanol for 45min, a mixture (v:v = 1:1) of ethanol and dimethylbenzene (Sinopharm, #10023418) for 8min, dimethylbenzene for 8min twice, and paraffin for 1h three times. 4-μm paraffin sections were processed for hematoxylin and eosin (H&E) staining or Sirius Red staining. For the immunohistochemistry, paraffin sections were dewaxed in dimethylbenzene for 15min three times, 100% ethanol for 5min twice, 85% ethanol for 5min, 75% ethanol for 5min, and sterile water for 5min twice. The sections were then incubated in citrate antigen retrieval solution at 95^°^C for 20min, followed by 3% (w/v) hydrogen peroxide solution for 25min. The sections were blocked with PBS / 3% BSA for 30min and sequentially immunostained with the primary rabbit anti-α-SMA antibody (Abcam, #ab5694) and the corresponding secondary HRP-conjugated goat anti-rabbit IgG antibody (Genetech, #GK500510A). The sections were then treated with DAB substrate (Solarbio, #DA1010) for 20min to visualize the immunohistochemistry signal and finally stained with hematoxylin. All the processed paraffin sections were scanned by Axio Scan Z1. The percentage of Sirius Red-positive area in each section was quantified in ImageJ. The anti-α-SMA immunohistochemistry signals were measured with the H-DAB model in the ImageJ plugin of Immunohistochemistry Image Analysis Toolbox (https://imagej.nih.gov/ij/plugins/ihc-toolbox/index.html).

### Quantitative PCR (qPCR) analysis

Liver tissues were freshly dissected from the mice of indicated conditions. Total RNAs were extracted by the RNeasy Mini Kit and analyzed by the SYBR Green Real-Time PCR Kit (Thermo Fisher Scientific). The primer sequences used for the qPCR analysis are *Col1a1* (forward: TTCTCCTGGCAAAGACGGAC; reverse: CGGCCACCATCTTGAGACTT), *Col1a2* (forward: CCCAGAGTGGAACAGCGATT; reverse: ATGAGTTCTTCGCTGGGGTG), *Col3a1* (forward: CATGCATAAATGCCAGCCCC; reverse: CCGGCTGGAAAGAAGTCTGA), *Timp1* (forward: TCTTGGTTCCCTGGCGTACTCT; reverse: GTGAGTGTCACTCTCCAGTTTGC), and *Esr1* (forward: TCTGCCAAGGAGACTCGCTACT; reverse: GGTGCATTGGTTTGTAGCTGGAC). *B2m* mRNA levels were utilized as the internal control (forward: CTCGGTGACCCTGGTCTTTC; reverse: GGATTTCAATGTGAGGCGGG).

### FACS and in vitro cultures of mouse HSCs

For the FACS sorting of mouse HSCs, the procedure was modified from the published protocol (Mederacke et al., 2015). The mice of indicated conditions were anesthetized and perfused via the inferior vena cava with 20 ml of the Perfusion Solution, followed by 30ml of the Digestion Solution containing 0.5mg/ml pronase and 40ml of the Digestion Solution containing 0.625mg/ml collagenase Ⅱ. The liver tissues were dissected out and further incubated in the Digestion Solution containing 0.625mg/ml collagenase Ⅱ and 75U/ml DNaseΙ at 37^°^C for 15 min. All the solutions were pre-warmed to 37^°^C before use. The liver samples were mashed through a 70-μm cell strainer and centrifuged at 50*g* for 3min at 4^°^C to remove tissue debris and hepatocytes, and the supernatant containing non-parenchymal cells was collected and further centrifuged at 580*g* for 10min at 4^°^C. The cell pellets were washed in GBSS and centrifuged again at 580*g* for 10min at 4^°^C. The cell pellets were resuspended in GBSS containing a final concentration of 10% (w/v) Nycodenz. 10ml of cell suspension was loaded into 15ml Falcon tubes and gently overlaid with 1.5 ml GBSS. After the centrifugation at 1,380*g* for 17min at 4^°^C, the resulting cell layers between the two fractions were collected. Those non-parenchymal cells of mouse liver tissues were stained with FITC-conjugated anti-mouse CD45 (BioLegend, #157608) and 7-AAD and sorted on BD Aria Fusion. Mouse HSCs were identified as retinol (405nm autofluorescence)^+^ CD45^-^, and the FACS data were processed by FlowJo (https://www.flowjo.com).

In vitro cultures of mouse HSCs were modified from the published methods (Arab et al., 2020; Mederacke et al., 2015). FACS-sorted HSCs from the liver tissues of C57BL/6 wild-type female mice were cultured in the Dulbecco’s Modified Eagle’s Medium (DMEM; Gibco, #C11995500BT) containing 10% fetal bovine serum (FBS; NewZerum, #FBS-UE500), 100 U/ml penicillin, and 100 μg/ml streptomycin for 2 days. Mouse HSCs were then changed to fresh medium without or with a final concentration of 10μM 17β-estradiol (Sigma, #E8875) and cultured for 4 days.

Total RNAs of mouse HSCs were extracted by the RNeasy Mini Kit and subjected to single-end RNA-seq by the Beijing Genomics Institute. Gene expression levels were normalized as transcripts per million (TPM). The bulk RNA-seq data was deposited to the Sequence Read Archive with the accession number GSE275942.

### Chromatin immunoprecipitation followed by sequencing (ChIP-seq)

*Lrat^Cre^; RCL-ChR2(H134R)/EYFP^+/-^* female mice were subjected to the model of liver fibrosis. HSCs identified as EYFP^+^ CD45^-^ were FACS-sorted from the mice following the procedure described above. 5 × 10^6^ HSCs were collected in 2ml DMEM containing 10% FBS, 100 U/ml penicillin, 100 μg/ml streptomycin, and 10μM 17β-estradiol. The cells were fixed with 1% formaldehyde at room temperature for 10min and then quenched with a final concentration of 0.3M glycine for 5min. The fixed cells were incubated in 1ml ice-chilled cell lysis buffer (10mM Tris, 10mM NaCl, 0.2% NP-40/Igepal, pH 8.0) supplemented with 1:500 protease inhibitor cocktail (Roche) and 1:100 phenylmethylsulfonyl fluoride (PMSF; Sigma) for 20min. After cell lysis, nuclei were pelleted at 1,000*g* for 2min at 4^°^C and then incubated in 1ml nuclear lysis buffer (50mM Tris, 10mM NaCl, 1% SDS, pH 8.0) supplemented with 1:500 protease inhibitor cocktail and 1:100 PMSF for 20min on ice. The nuclei lysate containing chromatin was sonicated on the Qsonica Q800R3 sonicator (80% amplitude, 20s ON/40s OFF for 17min). The sonicated samples were centrifuged at 15,000*g* for 10min at 4^°^C to remove cell debris, and chromatin in the supernatant was collected.

The chromatin samples were diluted with 4 volumes of Dilution Buffer (20mM Tris-HCl, 150mM NaCl, 2mM EDTA, 1% Triton X-100, 0.01% SDS, pH 8.0) supplemented with 1:500 protease inhibitor cocktail and 1:100 PMSF. 50μl of protein A/G agarose beads (Santa Cruz, #sc-2003) were added to each sample for pre-clearing at 4^°^C for 8h with gentle rotation. Each pre-cleared chromatin sample was then incubated with 35μl of protein A/G agarose beads pre-bound with 10μl of ChIP-grade rabbit anti-ESR1 antibody (Abcam, #ab32063) at 4℃ overnight. Protein A/G agarose beads were washed on ice once with Wash Buffer I (20mM Tris-HCl, 50mM NaCl, 2mM EDTA, 1% Triton X-100, 0.1% SDS, pH 8.0), twice with High Salt Buffer (20mM Tris-HCl, 500mM NaCl, 2mM EDTA, 1% Triton X-100, 0.01% SDS, pH 8.0), once with Wash Buffer II (10mM Tris-HCl, 250mM LiCl, 1mM EDTA, 1% NP-40, 1% sodium deoxycholate, pH 8.0), and twice with TE Buffer (10mM Tris-HCl, 1mM EDTA, pH 8.0). The immunoprecipitated chromatin was eluted from protein A/G agarose beads in 200µl Elution Buffer (100mM NaHCO_3_, 1% SDS) at room temperature and reverse-crosslinked in the presence of 30μg/ml proteinase K at 65℃ overnight. Each eluted sample was supplemented with 10μl of 3M sodium acetate (pH 5.2), and DNA was purified by the QIAquick PCR Purification Kit (Qiagen). The ChIP-seq libraries were constructed using the VAHTS Universal DNA Library Prep Kit for MGI (Vazyme, #NDM607-02) and sequenced on the MGI DNBSEQ-T7 platform. The obtained ChIP-seq data was deposited to the Sequence Read Archive with the accession number GSE275942.

The ChIP-seq reads were first filtered with Trim Galore (v0.6.6) and then aligned to mouse genome assembly mm9 using Bowtie2 (v2.3.5.1). Low-quality reads (MAPQ <30) were filtered out, and reads that aligned to mitochondria random contigs and ENCODE blacklisted regions were removed. In addition, PCR duplicates were removed using Picard (v2.23.3). Valid reads were then normalized using the counts per million (cpm) method through Deeptools (v3.1.3). Enrichment analysis of transcription factor motifs was performed on the narrow peak file using the findMotifsGenome.pl module of HOMER v.4.11 with the parameters “-size 200 -len 8 -mis 2 -S 5”. Background sequences were derived from the mm9 genome in the HOMER database. The top motifs from the known matches are visualized by the R package universal motif (v.1.18.1).

### Single-cell RNA sequencing (scRNA-seq) analyses

The published scRNA-seq datasets of the non-parenchymal cells of adult male mice, i.e., GSE136103 (Duan et al., 2021) and GSE137720 (Dobie et al., 2019), were obtained from the Sequence Read Archive. GSE136103 contains one dataset (GSM4041174) of the control healthy condition and one dataset (GSM4041175) of the model of CCl_4_-induced liver fibrosis. GSE137720 contains two datasets (GSM4085623 and GSM4085625) of the control healthy condition and three datasets (GSM4085624, GSM4085626, and GSM4085627) of the model of CCl_4_-induced liver fibrosis. For quality control, we filtered out the cells with <800 or >30000 unique molecular identifiers (UMIs), <500 or >6000 genes, or >10% mitochondrial reads. We further filtered out the doublets in each dataset through the DoubletFinder (https://github.com/chris-mcginnis-ucsf/DoubletFinder).

We loaded the filtered count matrixes to the CreateSeuratObject function in Seurat (v4.0) (https://github.com/satijalab/seurat) to create Seurat objects, which were then merged into one Seurat object, followed by the log-normalization by the NormalizeData function. The top 2,000 variable genes were identified using the FindVariableFeatures function. Principal component analysis (PCA) was performed using the RunPCA function, and batch effect correction was then conducted on the principal components with the Harmony function to integrate different datasets. Unsupervised clustering was performed using the FindNeighbors and FindClusters functions. For the determination of cell types, we referred to the cell identity information provided in the original publications (Dobie et al., 2019; Duan et al., 2021), and the feature plots of indicated genes were generated.

### Statistical methods

Student’s *t*-test (two-tailed unpaired) or ANOVA test (two-way with *post hoc* tests) was performed using GraphPad Prism 9.5.0 (http://www.graphpad.com/scientific-software/prism). All the data points in the figures represent biological replicates, and statistical details of the experiments are included in the figure legends.

## Acknowledgments

This work has been supported by the National Key Research and Development Program of China (#2023YFA1801900 to J.Y.), the National Natural Science Foundation of China (#32125017 to J.Y.; #82273104 and #82473149 to W.F.), and the Beijing Natural Science Foundation (#7232086 to J.Y.). Additional funds to J.Y.’s research group have been from the State Key Laboratory of Membrane Biology at Peking University and the Center for Life Sciences at Peking University.

## Author Contributions

J.Y. conceived and designed this study. T.L., H.Z., F.L., Z.D., Y.C., and H.Z. performed the experiments and analyzed the results. G.W., D.C., X.Z., and F.W. provided human liver tissues. T.L., Y.D., Y.C., and J.Y. prepared the manuscript.

